# Nutrient dynamics and stream order influence microbial community patterns along a 2914 km transect of the Mississippi River

**DOI:** 10.1101/091512

**Authors:** Michael W. Henson, Jordan Hanssen, Greg Spooner, Patrick Fleming, Markus Pukonen, Frederick Stahr, J. Cameron Thrash

**Author notes:** Correspondence: J. Cameron Thrash, Louisiana State University, Department of Biological Sciences, 202 Life Sciences Bldg., Baton Rouge, LA 70803.

## Abstract

Draining 31 states and roughly 3 million km^2^, the Mississippi River (MSR) and its tributaries constitute an essential resource to millions of people for clean drinking water, transportation, agriculture, and industry. Since the turn of the 20^th^ century, MSR water quality has continually rated poorly due to human activity. Acting as first responders, microorganisms can mitigate, exacerbate, and/or serve as predictors for water quality, yet we know little about their community structure or ecology at the whole river scale for large rivers. We collected both biological (16S and 18S rRNA gene amplicons) and physicochemical data from 38 MSR sites over nearly 3000 km from Minnesota to the Gulf of Mexico. Our results revealed a microbial community composed of similar taxa to other rivers but with unique trends in the relative abundance patterns among phyla, OTUs, and the core microbiome. Furthermore, we observed a separation in microbial communities that mirrored the transition from an 8^th^ to 10^th^ Strahler order river at the Missouri River confluence, marking a different start to the lower MSR than the historical distinction at the Ohio River confluence in Cairo, IL. Within MSR microbial assemblages we identified subgroups of OTUs from the phyla Acidobacteria, Bacteroidetes, Oomycetes, and Heterokonts that were associated with, and predictive of, the important eutrophication nutrients nitrate and phosphate. This study offers the most comprehensive view of MSR microbiota to date, provides important groundwork for higher resolution microbial studies of river perturbation, and identifies potential microbial indicators of river health related to eutrophication.

## Introduction

By connecting terrestrial, lotic, and marine systems, rivers perform vital roles in both the transport and processing of compounds in all major global biogeochemical cycles (Richey *et al.* 2002; Ensign and Doyle 2006; Withers and Jarvie 2008; Battin *et al.* 2009; Savio *et al.* 2015). Within the carbon cycle alone, rivers collectively discharge organic carbon to the oceans at over 0.4 Pg C yr^−1^ (Cauwet *et al.* 2002). Perhaps more importantly, rivers are generally net heterotrophic (Cole *et al.* 2007), indicating that they not only transport organic matter but host active metabolic processing of it as well. Conservative estimates place heterotrophic output of the world’s fluvial networks (streams, rivers, and estuaries) at 0.32 Pg C yr^−1^ (Cole and Caraco 2001; Battin *et al.* 2009). Although rivers contain a small minority of global fresh water at any given moment, the considerable volumes that pass through these systems make them relevant to models attempting to quantify global elemental transformations. However, despite the fact that microbial functions likely play a vital role in ecosystem health for both rivers themselves and their places of discharge, microbial functions in rivers remain understudied.

At 3734 km, the Mississippi River (MSR) is the fourth longest on earth, draining 31 U.S. states and two Canadian provinces- a watershed consisting of 41% of the continental U.S (Turner and Rabalais 2003; Dagg *et al.* 2004). The MSR is a major source of drinking water for many U.S. cities; a critical thoroughfare for transportation, commerce, industry, agriculture, and recreation; and conveys the vestiges of human activity to the Gulf of Mexico (GOM). In New Orleans, the average flow rate is over 16,990 cubic meters s^−1^ (cms) (Rabalais *et al.* 1996), but can exceed 84,000 cms during flood stages (Singh 2012), and carries over 150 × 10^9^ kg of suspended sediment into the northern GOM annually (Dagg *et al.* 2004, 2005). The MSR also transports considerable amounts of carbon, nitrogen and phosphorus, with average annual fluxes of 2.1 TgC yr^−1^, > 1.4 TgN year^−1^, and > 0.14 TgP yr^−1^, respectively (Goolsby and Battaglin 2001; Cauwet *et al.* 2002; Aulenbach *et al.* 2007). Globally, the MSR represents 0.8% of the dissolved organic carbon flux to the worlds oceans (Cauwet *et al.* 2002). When considering total nitrogen, up to 62% can occur as nitrate (NO_3_^−^) (Goolsby and Battaglin 2001). This massive discharge of excess eutrophic nutrients, primarily from corn and soybean (nitrogen) and animal manure (phosphorus) runoff (McIsaac *et al.* 2001; Turner and Rabalais 2004; Alexander *et al.* 2008; Schilling *et al.* 2010; Duan *et al.* 2014; Staley *et al.* 2014a), fuels one of the largest marine zones of seasonal coastal hypoxia in the world (Rabalais *et al.* 2002, 2007; Bianchi *et al.* 2010; Bristow *et al.* 2015). Studying microbial relationships to river eutrophication will improve our understanding of their contributions to either mitigating or exacerbating nutrient input.

Far from a homogenous jumble of organisms ferried downriver, microbial community composition changes with distance from the river mouth and/or from the influence of tributaries (Kolmakova *et al.* 2014; Read *et al.* 2015; Savio *et al.* 2015), resulting from altered nutrient concentrations (Staley *et al.* 2014a; Van Rossum *et al.* 2015; Meziti *et al.* 2016), differing dissolved organic matter (DOM) sources (Ruiz-González *et al.* 2013; Zeglin 2015; Blanchet *et al.* 2016), and land use changes (Staley *et al.* 2014b; Van Rossum *et al.* 2015; Zeglin 2015). Past studies of the Thames, Danube, Yenisei, and Columbia Rivers have found that planktonic river microbial assemblages were dominated by the phyla Actinobacteria, Proteobacteria, and Bacteriodetes; and taxa such as acI Actinobacteria, *Polynucleobacter* spp., GKS9 and LD28 *Betaproteobacteria*, CL500-29 Actinobacteria, LD12 SAR11 *Alphaproteobacteria*, and *Novosphingobium* spp. (Crump *et al.* 1999; Kolmakova *et al.* 2014; Read *et al.* 2015; Savio *et al.* 2015).

Specific to the MSR, previous 16S rRNA gene amplicon and metagenomic studies have demonstrated that microbial assemblages in the Minnesota portion of the river (Lake Itasca to La Crescent) correlated with organic carbon concentration, total dissolved solids, and land use changes (Staley *et al.* 2013, 2014a; b). Species richness increased near developed land with greater concentrations of nitrate and nitrite, as opposed to sites near pastureland that had greater organic carbon (Staley *et al.* 2014a). Similar to past studies of other lotic systems (Crump *et al.* 2012; Read *et al.* 2015; Savio *et al.* 2015), Actinobacteria and Proteobacteria increased in relative abundance along the upper MSR, while OTUs belonging to Bacteroidetes decreased (Staley *et al.* 2013). A follow up time-series study over two summers showed a distinct seasonal signal: samples from late summer of both years were more similar to each other than those from early summer (Staley *et al.* 2015). A more recent study of another portion of the MSR (above the Missouri River confluence to Natchez, La.) found bacterial communities formed distinct regimes, likely due to the strong influence of their respective tributaries (Jackson *et al.* 2014; Payne *et al.* 2017).

Researchers have suggested that some of these patterns supported the application of the River Continuum Concept (RCC) (Vannote *et al.*, 1980) to river microbiota. The RCC postulates that as a river increases in size, the influences of riparian and other inputs will decrease as the river establishes a dominant core community (Vannote *et al.* 1980). Richness will increase with stream order complexity before decreasing in higher order rivers (Vannote *et al.*, 1980). Therefore, as continuous systems with increasing volumes and residence times, river microbiota should transition from experiencing strong influences of mass effects from terrestrial, riverbed, and tributary sources to systems where species sorting plays a more important role (Crump *et al.* 2012; Besemer *et al.* 2013; Savio *et al.* 2015; Niño-García *et al.* 2016).

Accordingly, studies examining the linkages between RCC and microbial structuring mechanisms (e.g. species sorting, mass effects) have found that some of the results indeed supported the RCC. Specifically, numerous lotic studies found that upstream and headwaters amassed high species richness, before richness decreased further downstream (Crump *et al.* 2012; Kolmakova *et al.* 2014; Savio *et al.* 2015; Niño-García *et al.* 2016). This follows the expected pattern outlined by the RCC where early stream and river microbial communities are heavily influenced by a large input of allocthonous organisms from sediment and groundwater into the river network. As the rivers progressed, such mass effects likely heeded to species sorting, a process requiring an increased water residence time to allow for the selection of species based on local environmental conditions (Crump *et al.* 2012; Savio *et al.* 2015; Niño-García *et al.* 2016). These types of river modulations, influenced by hydrology and environmental conditions, have correlated with decreased species richness and increased core community membership (the taxa found consistently throughout the entire river) in multiple systems (Crump *et al.* 2012; Besemer *et al.* 2013; Savio *et al.* 2015; Niño-García *et al.* 2016). An analyses of an upper portion of the MSR provided evidence for the importance of species sorting (Staley *et al.* 2015). However, in a study of the Thames River, researchers found no association between microbial assemblages and any measured variables other than residence time, leading them to suggest that ecological succession was the primary driver of microbial community composition (Read *et al.* 2015). Furthermore, a 1300 km transect of the lower MSR indicated that richness actually increased instead of decreasing in particle-associated, and to a lesser extent, free-living, communities, suggesting that the MSR may have fundamentally different mechanisms driving microbial diversity than other rivers (Payne *et al.* 2017).

Complicating matters, particle-associated communities in rivers (frequently defined as those found on filters of > ~3 μm remain distinct from their free-living counterparts (Crump *et al.* 1999; Riemann and Winding 2001; Jackson *et al.* 2014), potentially due to increased production rates from access to readily obtainable carbon (Crump and Baross 1996; Crump *et al.* 1998, 1999). Typical river microorganisms associated with particles include OTUs related to the Bacteroidetes clades *Flavobacteria* and *Cytophaga*, Planctomycetes, *Rhizobium* spp., and *Methylophilaceae* spp. (Crump and Baross 1996; Allgaier and Grossart 2006; D’Ambrosio *et al.* 2014; Jackson *et al.* 2014). However, denoting consistent trends in particle community composition must be tempered by recent evidence suggesting substrate availability and chemical queues may trigger organisms to switch between free-living and particle-associated lifestyles (Grossart 2010; D’Ambrosio *et al.* 2014). Rivers constitute complex and highly dynamic ecosystems from a metacommunity perspective.

Although important insights have been gained from recent research on portions of the MSR (Staley *et al.* 2013, 2015, 2016; Payne *et al.* 2017), microbiological transects at the whole-river or catchment scale have yet to be completed. This study aimed to *i)* compare the structure and abundance of size-fractionated MSR microbial community populations to those in other rivers, *ii)* examine within-river heterogeneity of microbial communities, and *iii)* identify MSR microorganisms most strongly associated with eutrophication- all at a near whole-river scale, for both prokaryotes and microscopic eukaryotes. During the fall of 2014, we completed the most extensive microbiological survey of the Mississippi River to date with a continual rowed transect over 70 days. Rowers from the adventure education non-profit OAR Northwest collected samples from Minneapolis, MN to the Gulf of Mexico (2918 km) (Fig. 1A). Our findings expand the current information available on microbial assemblages in major lotic ecosystems; further delineate the relationships between microbial structure and stream order, nutrients, and volume; and identify MSR taxa predictive of the eutrophication nutrients nitrate and phosphate.

**Figure 1.**
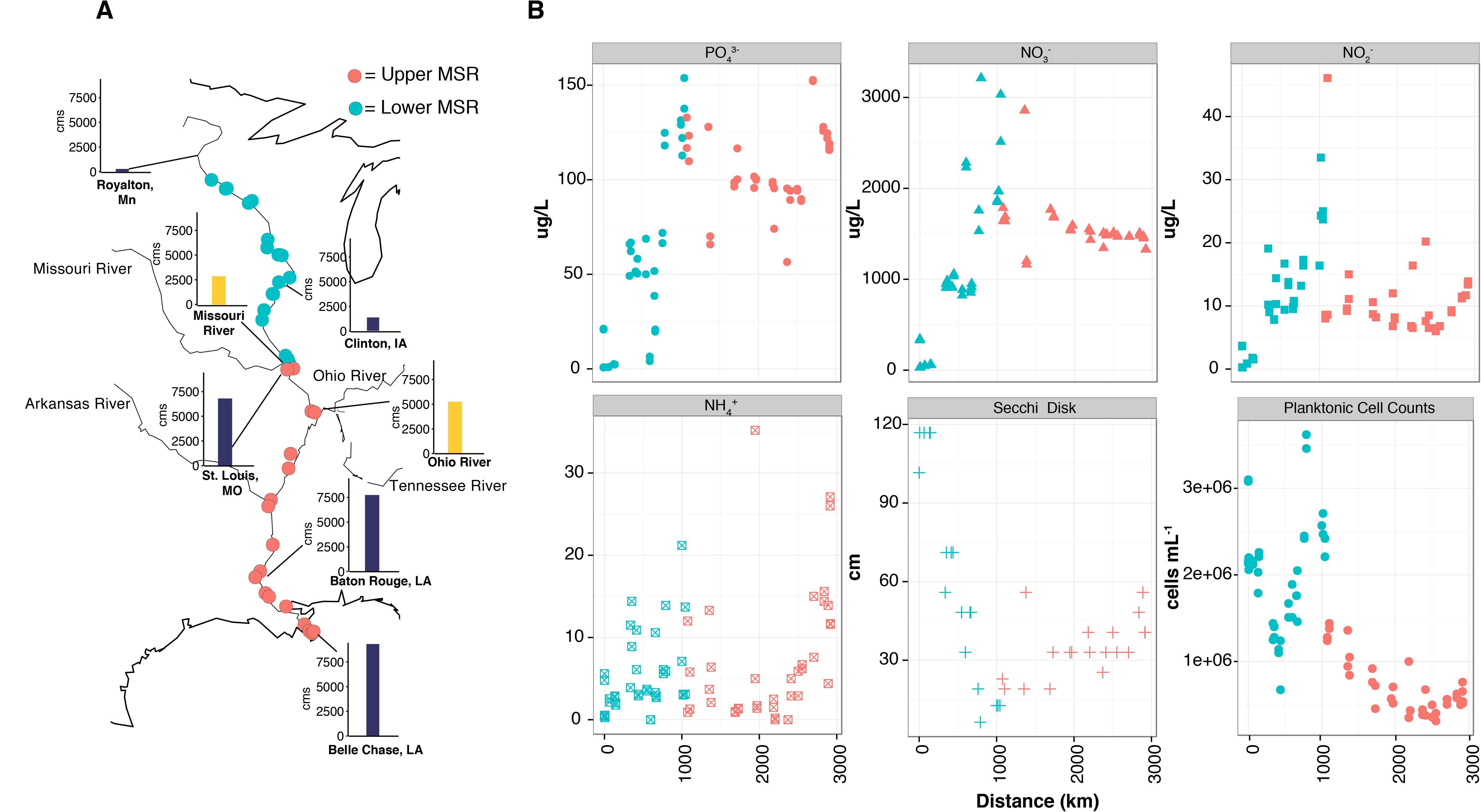
Sampling map (A) with graph inserts that represent the measured discharge rate (cubic meter second^−1^ [cms]) as recorded on the USGS gauges, and six environmental parameters measured along the transect, according to concentration or visible depth of secchi disk by distance (B). Throughout the figure and text, blue and red dots represent sampling locations above and below the Missouri River confluence, respectively, and are designated throughout as “upper” and “lower.” Cell Counts only represent the Planktonic (< 2.7 μm) fraction.

## Materials and Methods

### Sampling and Cell Counts

We used rowboats and a simple filtration protocol (Supplementary Information) to collect water from 39 sites along a continual-rowed transect of the MSR, starting in Lake Itasca and ending in the GOM, over 70 days from September 18^th^ to November 26^th^, 2014. Sites were chosen to be near major cities and above and below large tributaries. After some samples were removed due to insufficient sequence data, contamination, or incomplete metadata (see below), the final usable set of samples included 38 sites starting at Minneapolis (Fig. 1A, Table S1, General Information). Most sampling occurred within the body of the river, although due to safety issues, three samples were collected from shore (Table S1, General Information). We collected duplicate samples at each site, but because separate rowboat teams frequently collected these sometimes several dozen meters apart, they cannot be considered true biological replicates and we have treated them as independent samples.

At each site, we filtered 120 mL of water sequentially through a 2.7 μm GF/D filter (Whatman GE, New Jersey, USA) housed in a 25 mm polycarbonate holder (TISCH, Ohio, USA) followed by a 0.2 μm Sterivex filter (EMD Millipore, Darmstadt, Germany) with a sterile 60 mL syringe (BD, New Jersey, USA). We refer to fractions collected on the 2.7 μm and 0.22 μm filters as > 2.7 μm and 0.2-2.7 μm, respectively. Flow-through water from the first 60 mL was collected in autoclaved acid-washed 60 mL polycarbonate bottles. Both filters were wrapped in parafilm, and together with the filtrate, placed on ice in Yeti Roadie 20 coolers (Yeti, Austin, TX) until shipment to LSU. Further, 9 mL of whole water for cell counts was added to sterile 15 mL Falcon tubes containing 1 mL of formaldehyde and placed into the cooler. The final cooler containing samples from sites P-Al had substantial ice-melt. Though our filters were wrapped in parafilm, we processed melted cooler water alongside our other samples to control for potential contamination in these filters. Given that some of our samples were expected to contain low biomass, we also included duplicate process controls for kit contamination (Salter *et al.* 2014; Weiss *et al.* 2014) with unused sterile filters.

Flow-through 0.2 μm filtered water from each collection was analyzed for SiO_4_, PO_4_^3−^, NH_4_+, NO_3_^−^, and NO_2_^−^ (μg/L) at the University of Washington Marine Chemistry Laboratory (http://www.ocean.washington.edu/story/Marine+Chemistry+Laboratory). Aboard-rowboat measurements were taken for temperature and turbidity. We determined turbidity by deploying a secchi disk (Wildco, Yulee, FL), while drifting with the current so the line hung vertically. It was lowered until no longer visible, then raised until just visible, and measured for its distance below the waterline. We then calculated secchi depth from the average of two measurements. Temperature was measured with probes from US Water Systems (Indianapolis, IN), rinsed with distilled water between samples. Samples for cell counts were filtered through a 2.7 μm GF/D filter, stained with 1x Sybr Green (Lonza), and planktonic cells were enumerated using the Guava EasyCyte (Millipore) flow cytometer as previously described (Thrash *et al.* 2015).

Throughout the transect, seven cooler exchanges occurred to ensure samples were not exposed to extended durations at *in situ* temperatures. Cooler temperatures were monitored with HOBO loggers (Onset, Bourne, MA) to ensure that samples stayed at ≤ 4°C. On average, a sample spent 7.6 days (median = 6 days, range=1 day minimum, 16 days maximum) at ≤ 4°C before delivery to LSU. Though samples were not frozen immediately, previous research has provided evidence that soil and water samples stored at 4°C, and for different durations (≥ 14 days), did not show significant alterations to most community relative abundances, structure, or composition (Lauber *et al.* 2010; Rubin *et al.* 2013; Tatangelo *et al.* 2014).

### DNA extraction and Sequencing

DNA was extracted from both filter fractions and controls using a MoBio PowerWater DNA kit (MoBio Laboratories, Carlsbad, CA) following the manufacturer’s protocol with one minor modification: in a biosafety cabinet (The Baker Company, Stanford, ME), Sterivex filter housings were cracked open using sterilized pliers and filters were then removed by cutting along the edge of the plastic holder with a sterile razor blade before being placed into bead-beating tubes. DNA was eluted with sterile MilliQ water, quantified using the Qubit2.0 Fluorometer (Life Technologies, Carlsbad, CA), and stored at −20° C. Bacterial and archaeal sequences were amplified at the V4 region of the 16S rRNA gene using the 515f and 806r primer set (Caporaso *et al.* 2012), and eukaryotic sequences from the V9 region of the 18S rRNA gene using the 1391r and EukBR primer set (Amaral-Zettler *et al.* 2009). Amplicons were sequenced on an Illumina MiSeq as paired-end 250 bp reads at Argonne National Laboratory. Sequencing of the 16S and 18S rRNA gene amplicons resulted in 13,253,140 and 13,240,531 raw sequences, respectively.

### Sequence Analysis

We analyzed amplicon data with Mothur v.1.33.3 (Schloss *et al.* 2009) using the Silva v.119 database (Pruesse *et al.* 2007; Quast *et al.* 2013). Briefly, 16S and 18S rRNA gene sequences were assembled into contigs and discarded if the contig had any ambiguous base pairs, possessed repeats greater than 8 bp, or were greater than 253 or 184 bp in length, respectively. Contigs were aligned using the Silva rRNA v.119 database, checked for chimeras using UCHIME (Edgar *et al.* 2011), and classified also using the Silva rRNA v.119 database. Contigs classified as chloroplast, eukaryotes, mitochondria, or “unknown;” or as chloroplast, bacteria, archaea, mitochondria, or “unknown;” were removed from 16S or 18S rRNA gene data, respectively. The remaining contigs were clustered using the cluster.split() command into operational taxonomic units (OTUs) using a 0.03 dissimilarity threshold (OTU_003_). After these steps, 146,725 and 131,352 OTUs remained for the 16S and 18S rRNA gene communities, respectively.

### Sample quality control

To evaluate the potential for contamination from extraction kits, cooler water in the last set of samples, or leaking/bursting of pre-filters, all samples were evaluated with hierarchical clustering and NMDS analysis. Hierarchical clustering was performed in R using the *hclust* function with methods set to “average”, from the vegan package (Oksanen *et al.* 2015). Samples were removed from our analysis if they were observed to be outliers in both the NMDS and hierarchical clustering such that they grouped with our process controls. The process and cooler water controls were extreme outliers in both, as was sample L2 (Fig. S1, S2). Sterivex and prefilter samples generally showed strong separation with the exception of three 16S rRNA gene samples- STER X2, W2, S2 (Fig. S1, S2). The only other samples that were removed were due to missing chemical data (Lake Itasca1-2, A1-2) or failed sequencing (16S STER Af1; 16S PRE S2, X2; 18S PRE O1). Not including process or cooler water controls, 152 samples were sequenced each for prokaryotic and eukaryotic communities. After these QC measures, 144 and 149 samples remained in the analyses from the 16S and 18S rRNA gene amplicons, respectively. Further, to control for potential contaminants, any OTU with greater 20 reads in the process or cooler controls was removed from the data set. 146,725 and 131,327 OTUs remained after these steps for 16S and 18S rRNA gene communities, respectively.

### Alpha and Beta Diversity

OTUo.o3 analyses were completed with the R statistical environment v.3.2.1(R Developement Core Team 2015). Using the package PhyloSeq (McMurdie and Holmes 2013), alpha-diversity was first calculated on the non-normalized, unfiltered OTUs using the “estimate richness” command within PhyloSeq, which calculates six alpha diversity metrics (McMurdie and Holmes 2013). After estimating alpha diversity, potentially erroneous rare OTUs, defined here as those without at least two sequences in 20% of the sites, were discarded. After this filter, the dataset contained 950 and 724 16S and 18S rRNA gene OTUs, respectively. For site-specific community comparisons, OTU counts were normalized using the package DESeq2 (Love *et al.* 2014) with a variance stabilizing transformation (Learman *et al.* 2016). Beta-diversity between samples was examined using Bray-Curtis distances via ordination with non-metric multidimensional scaling (NMDS). Analysis of similarity (ANOSIM) was used to test for significant differences between groups of samples (e.g lower versus upper MSR) using the *anosim* function in the vegan package (Oksanen *et al.* 2015). The influence of Hellinger transformed environmental parameters on beta-diversity was calculated in R with the *envfit* function. Relative abundances plots were made using the R package ggplot2. Linear and nonlinear regressions were made using the command *stat_smooth()* with the flag “level=0.95” and method = “lm” or “loess” for linear and non-linear regressions, respectively.

### Network analyses and modeling

To identify specific OTUs with strong relationships to environmental parameters (e.g. turbidity, NO3”), we employed weighted gene co-expression network analysis (WGCNA) (Langfelder and Horvath 2008) as previously described for OTU relative abundances (Guidi *et al.* 2016). First, a similarity matrix of nodes (OTUs) was created based on pairwise Pearson correlations across samples. This was transformed into an adjacency matrix, which represents the strength at which two OTUs within the matrix are connected (adjacency). To do this, the similarity matrix was raised to a soft threshold power (p; p = 6 for 16S and 18S > 2.7 μm, p = 4 for 16S 0.2-2.7 μm, p = 9 for 18S 0.2-2.7 μm) that maximizes OTU connections with others and ensures scale-free topology fit. Submodules of highly co-correlating OTUs were assigned a color (e.g. yellow, green, turquoise) as defined with a topological overlap measure and hierarchical clustering, which identified submodules based on an OTU’s weighted correlations to other OTUs within and outside the network. Each submodule, represented by an eigenvalue, was pairwise Pearson correlated to individual Hellinger-transformed environmental measurements (Figs. S7-10A). To explore the relationship of submodule structure to these parameters, OTUs within the submodule were plotted using their individual correlation to the parameter of interest (y-axis, here nitrate or phosphate) and their submodule membership (x-axis), defined as the strength of an OTU (number of connections) to other OTUs within the submodule (Figs. S7-10B, D). Strong correlations between submodule structure and an environmental parameter facilitate identification of OTUs that are highly correlated to that parameter. To evaluate the predictive relationship between a submodule and a parameter, we employed partial least square regression (PLS) analysis. PLS maximizes the covariance between two parameters (e.g., OTU abundances and nitrate concentration) to define the degree to which the measured value (OTU abundance) can predict the response variable (nutrient concentration). The PLS model was permutated 1000 times and Pearson correlations were calculated between the response variable and leave-one-out cross-validation (LOOCV) predicted values. Modeled values were then compared with measured values to determine the explanatory power of the relationships (Figs. S7-10C, E). Relative contributions of individual OTUs to the PLS regression were calculated using value of importance in the projection (VIP) (Chong and Jun 2005) determination. PLS was run using the R package *pls* (Mevik and Wehrens 2007), while VIP was run using additional code found here: http://mevik.net/work/software/VIP.R. Though other environmental parameters had strong Pearson correlations to submodules, only submodules with strong, positive correlations to measures related to eutrophication were selected, in accordance with our study objectives, as these were most likely to provide candidate taxa with remediation potential.

### Accession numbers

Community 16S and 18S rRNA gene sequence fastq files are available at the NCBI Sequence Read Archive under the accession numbers: SRR3485674 - SRR3485971 and SRR3488881 - SRR3489315, respectively.

### Code Availability

All code used for Mothur, SeqENV, PhyloSeq, WGCNA, and PLS regression analyses can be found on the Thrash lab website (http://thethrashlab.com/publications) with the reference to this manuscript linked to “Supplemental Information”.

## Results

Using rowboats and a simple syringe-based filtration protocol, we measured 12 biological, chemical, and physical parameters (e.g. 16S and 18S rRNA gene communities, NH_4_+, river speed, etc.) from 38 sites along a 2918 km transect of the MSR (Fig. 1A). River order increases dramatically at the Missouri confluence [eighth to tenth Strahler order (Pierson *et al.* 2008)], which corresponded to overall discharge (Fig. 1A) and beta diversity changes discussed below. Therefore, we refer to this juncture as the separator between the upper (0 km - 1042 km, Sites AS) and lower MSR (1075-2914 km, Sites T-Al). Within the upper MSR, NO_3_^−^, PO_4_^3−^, and NO_2_^−^ were variable but generally increased downriver until peak concentrations near the confluences of the Illinois and Missouri Rivers. This gave way to lower and more consistent concentrations along the lower MSR (Fig. 1B). Turbidity (inversely related to secchi disk visibility) increased steadily downriver to a maximum near the Illinois and Missouri River confluences (1042 km, Site S) (Fig. 1B, Secchi Disk), then trended downwards for the rest of the transect. Planktonic (< 2.7 μm) cell counts varied between 1 and 3×10^6^ cells ·mL^−1^ in the upper MSR, and decreased to high 10^5^ cells · mL^−1^ in the lower MSR (Fig. 1B, Planktonic Cell Counts). Water temperature ranged from 19°C (133km, Site E) to 11.7°C (2552 km, Site Ag), and river speed, excluding three sites sampled from shore, was between 5.5 mph at Site Y and 0.4 mph (597 km, Site L) (Table S1, General Information). Spearman rank correlations of the measured environmental parameters showed strong positive correlations between nitrate, phosphate, distance, and increased turbidity; while nitrate and phosphate both strongly correlated negatively to water temperature and river speed (Table S1, Spearman Rank).

### Bacterial and archaeal communities

We observed a clear distinction between the 0.2-2.7 μm and > 2.7 μm 16S rRNA gene communities (ANOSIM R = 0.65, P = 0.001) (Fig. S1A). Both size fractions also showed significant community separation between sites above and below the Missouri River confluence (ANOSIM, > 2.7 μm: R = 0.44, P=0.001; 0.2-2.7 μm: R = 0.48, P = 0.001) (Fig. 2A, B), that the Eukaryotic fractions mirrored (below)(Fig. 2C, D). Overall richness was significantly higher in the lower vs. upper rivers (Table S1, Richness comparisons). Taken separately, most measurements indicated that richness increased with river distance in the upper and lower rivers, and this trend was common for both the > 2.7 μm and 0.2-2.7 μm fractions (Fig. S3A, B). Evenness measurements had less consistent trends.

**Figure 2.**
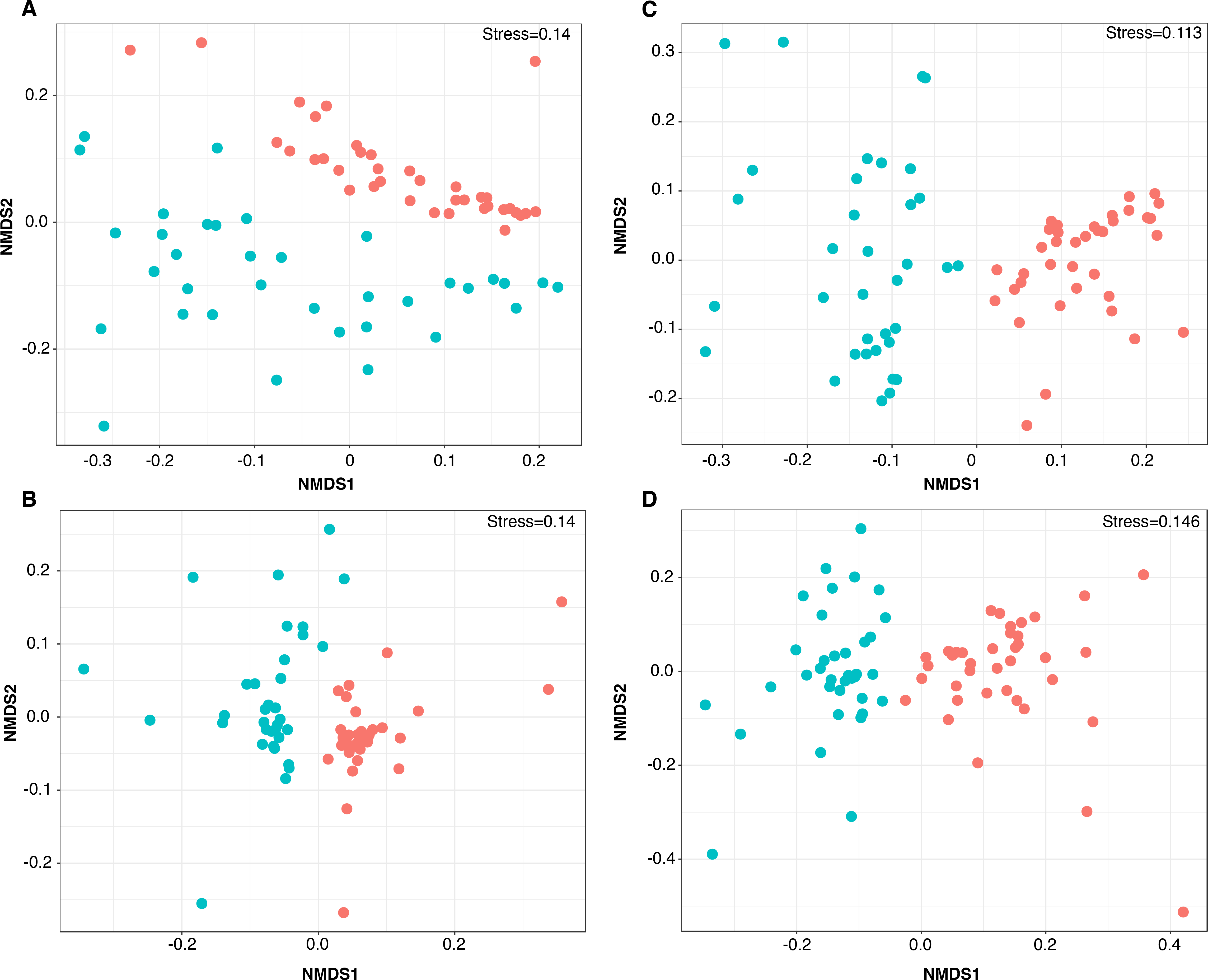
Non-metric multidimensional scaling (NMDS) results for whole community. The four plots represent > 2.7 μm (A, C) and 0.2-2.7 μm (B, D) fractions for the 16S (A, B) and 18S (C, D) rRNA gene communities.

Phosphate and turbidity had the highest correlation with the separation between the upper and lower > 2.7 μm communities (R^2^ = 0.58, R^2^ = 0.54, respectively), with water temperature (R^2^ = 0.52) also contributing (Table 1). At an OTU level, taxa related to the acI clade (Actinobacteria) and unclassified *Bacillaceae* (R^2^ > 0.77, P = 0.001) contributed most to the separation between the upper and lower > 2.7 communities, with OTUs related to the *Bacillales*, *Gemmatimonadaceae*, *Peptococcaceae*, and *Micromonosporaceae* clades also a factor (R^2^ > 0.70, P = 0.001) (Table 2). For the 0.2-2.7 μm fraction, nitrate were the strongest correlating environmental factors with the distinction between upper and lower communities (R^2^ = 0.443), although phosphate, turbidity, and water temperature (R^2^ > 0.385 for each) also contributed (Table 1). OTUs related to *Flavobacterium* and an unclassified Bacterium (closest NCBI BLAST hit *Acidovorax* sp., KM047473), most strongly contributed to the separation between the 0.2-2.7 μm communities (R^2^ > 0.503, P = 0.0001) (Table 2). Other important OTUs driving the separation between upper and lower communities belonged to the clades Bacteroidetes, *Microbacteriaceae*, *Clostridiales*, and *Holophagaceae* (R^2^ > 0.49, P = 0.0001) (Table 2).

**Table 1.** Summary of the correlations between environmental variables and microbial community beta diversity ordination on NMDS plots in Figure 2. +/-designates the direction of the association between the environmental variab e and NMDS axis

| Variable |  | NMDS1 | NMDS2 | P-value | R<sup>2</sup> |  | NMDS1 | NMDS2 | P-value | R<sup>2</sup> |
| --- | --- | --- | --- | --- | --- | --- | --- | --- | --- | --- |
| *PO<sub>4</sub><sup>3-</sup>* | 16S rRNA Gene 0.2-2.7 μm | + | + | 0.001 | 0.404 | 16S rRNA Gene > 2.7 μm | + | + | 0.001 | 0.575 |
| *SiSO<sub>4</sub>* |  | - | + | 0.003 | 0.183 |  | - | - | 0.015 | 0.115 |
| *NO<sub>3</sub><sup>-</sup>* |  | + | + | 0.001 | 0.443 |  | + | + | 0.001 | 0.33 |
| *NH<sub>4</sub><sup>+</sup>* |  | - | - | 0.002 | 0.178 |  | - | - | 0.004 | 0.189 |
| *NO<sub>2</sub><sup>-</sup>* |  | Not significant |  |  |  |  | Not significant |  |  |  |
| *River Speed* |  | - | - | 0.001 | 0.261 |  | - | - | 0.001 | 0.39 |
| *Water Temperature* |  | - | - | 0.001 | 0.385 |  | - | - | 0.001 | 0.524 |
| *Turbidity* |  | - | - | 0.001 | 0.409 |  | - | - | 0.001 | 0.536 |
| *PO<sub>4</sub><sup>3-</sup>* | 18S rRNA Gene 0.2-2.7 μm | + | + | 0.001 | 0.555 | 18S rRNA Gene > 2.7 μm | + | - | 0.001 | 0.486 |
| *SiSO<sub>4</sub>* |  | - | + | 0.001 | 0.264 |  | - | - | 0.001 | 0.265 |
| *NO<sub>3</sub><sup>-</sup>* |  | Not significant |  |  |  |  | + | + | 0.001 | 0.2 |
| *NH<sub>4</sub><sup>+</sup>* |  | - | + | 0.019 | 0.106 |  | Not significant |  |  |  |
| *NO<sub>2</sub><sup>-</sup>* |  | Not significant |  |  |  |  | Not significant |  |  |  |
| *River Speed* |  | - | - | 0.001 | 0.282 |  | - | - | 0.001 | 0.345 |
| *Water Temperature* |  | - | - | 0.001 | 0.29 |  | - | - | 0.001 | 0.372 |
| *Turbidity* |  | - | - | 0.001 | 0.272 |  | - | - | 0.001 | 0.353 |

**Table 2.**
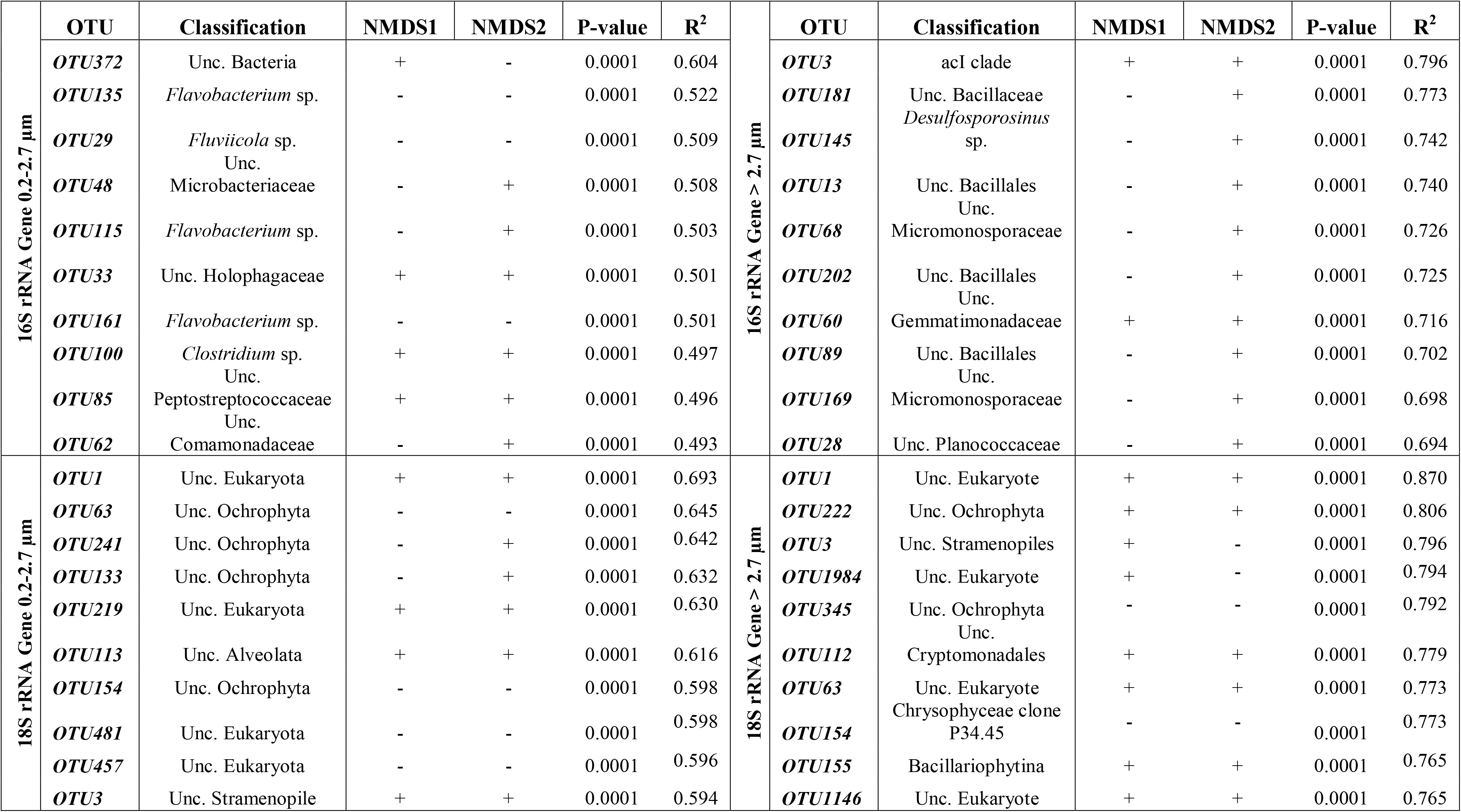
Summary of the correlations between OTUs and microbial community beta diversity ordination on NMDS plots in Figure 2. +/- designates the direction of the association between the OTU and NMDS axis.

At the phylum level, Proteobacteria, Actinobacteria, and Bacteroidetes dominated bacterial communities in both fractions (Figs. 3A and B) (Table S1, 16S OTU table norm.). Proteobacteria abundance in the > 2.7 μm fraction fluctuated over the course of the transect (Fig. 3A), whereas their 0.2-2.7 μm counterparts generally increased in relative abundance downriver (Fig. 3B). 0.2-2.7 μm Bacteroidetes and Actinobacteria generally decreased in the upper river and stabilized in the lower river. These phyla showed considerable abundance variation in the > 2.7 μm fraction. Cyanobacteria in the > 2.7 μm fraction negatively correlated with increased turbidity (Spearman rank = 0.67), consistent with lower irradiance (Fig. 3A). Both > 2.7 μm and 0.2-2.7 μm Acidobacteria increased in abundance downriver and positively correlated with river distance (Wilcoxon single ranked test, P = < 0.01) (Fig. S4A, B). Within the 0.2-2.7 μm fraction, the five most abundant OTUs were classified as a LD12 (OTU11; Proteobacteria), two acI clade OTUs (OTU4, OTU7; Actionbacteria), a *Limnohabitans* sp. (OTU2; Proteobacteria), and a LD28 (OTU8; Proteobacteria) (Table S1, 16S OTU table norm.). Comparatively, an unclassified *Methylophilaceae* (OTU1; Proteobacteria), a *Planktothrix* sp. (OTU12; Cyanobacteria), a NS11-12 marine group (OTU21; Bacteroidetes), an Candidatus Aquirestis sp. (OTU17; Bacteroidetes), and an unclassified *Sphingobacteriales* (OTU25; Bacteroidetes) were the most abundant OTUs in the > 2.7 μm fraction (Table S1, 16S OTU table norm.). Archaea occurred at much lower relative abundances: we found only eight OTUs belonging to the phyla Euryarchaeota and Thaumarchaeota, collectively. These OTUs were classified as *Methanobacterium* sp. (OTU714, 1968) and soil Crenarchaeotic Group (OTU370, 389, 983, and 1253), *Candidatus* Nitrosoarchaeum sp. (OTU1093) from the phylum Euryarcheota, and an unclassified Thaumarchaeota (OTU2951) from the phylum Thaumarchaeota (Table S1, 16S OTU table norm.). Overall, Thaumarchaeota increased in abundance along the transect more in the > 2.7 μm fraction compared to the 0.2-2.7 μm fraction (Fig. S4A, B). In both fractions, we only detected Euryarcheota at very low abundances. Importantly, the primers used in this study may miss some archaeal taxa (Parada *et al.* 2015).

**Figure 3.**
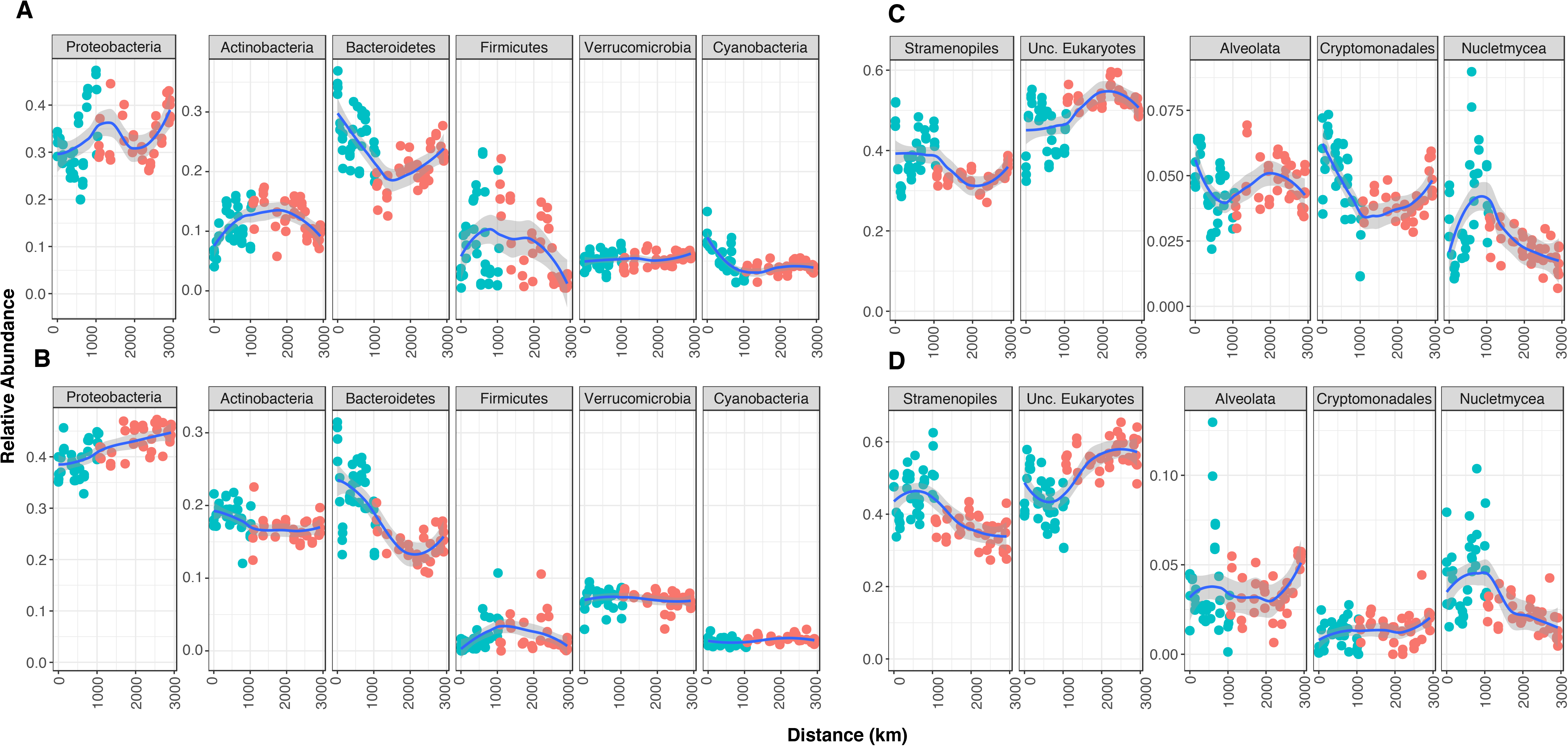
Relative abundance, according to transect distance, for the top 6 and 5 phyla in the 16S rRNA gene > 2.7 μm (A) and 0.2-2.7 μm (B) communities and 18S rRNA gene > 2.7 μm (C) and 0.2-2.7 μm (D) communities, respectively. Non-linear regressions with 95% CI (gray shading) are provided for reference.

### Microbial eukaryotic communities

Eukaryotic communities, observed via the 18S rRNA gene, also showed a significant separation between > 2.7 μm and 0.2-2.7 μm fractions (ANOSIM R = 0.689, P = 0.001) (Fig. S1B). As expected due to generally larger cell sizes in microbial eukaryotes compared to prokaryotes, species richness remained higher in the > 2.7 μm vs. 0.2-2.7 μm fractions (Fig. S3C, D). Both the > 2.7 μm and 0.2-2.7 fractions also showed a significant separation between the upper and lower MSR communities (ANOSIM, > 2.7 μm: R = 0.696, P = 0.001; 0.2-2.7 μm: R = 0.576, P = 0.001) (Fig. 2C, D). Overall richness in the > 2.7 μm fraction was also higher in the lower vs. upper river, but this was not true for the 0.2-2.7 μm fraction (Table S1, Richness comparisons). Richness in the > 2.7 μm fraction increased along both the upper and lower river, similarly to prokaryotic communities, but remained relatively stable within the 0.2-2.7 μm fraction.

Phosphate was the top environmental factor correlating to the distinction between eukaryotic communities in both filter fractions (> 2.7 μm, R^2^ = 0.49; 0.2-2.7 μm R^2^ = 0.56) (Table 1). No other factors had correlations > 0.38 (Table 1). At the OTU level, taxa related to an unclassified Ochrophyta (OTU63) and an unclassified Eukaryote (OTU1) separated the 0.2-2.7 μm communities (R^2^ > 0.645, P = 0.0001), while the same unclassified Eukaryote OTU (OTU1) and a second unclassified Eukaryote (OTU222) contributed most to separating the > 2.7 μm communities (R^2^ > 0.80, P = 0.0001) (Table 2).

Stramenopiles (or Heterokonts), encompassing diatoms and many other forms of algae, and OTUs that could not be classified at the phylum level dominated both the > 2.7 μm and 0.2-2.7 μm communities (Fig. 3C, D). Stramenopiles accounted for over 25% of both communities, with higher abundances in the upper vs. lower river. We observed a similar trend of disparate abundances between the upper and lower river for > 2.7 μm Cryptomonadales and 0.2-2.7 μm Nucletmycea, the latter of which include fungi (Fig. 3C; Table S1, 18S OTU table norm.). Within the 0.2-2.7 μm fraction, we identified the five most abundant OTUs as three unclassified *Bacillariophytina* (OTU7, OTU14, OTU9), a *Pythium* sp. (OTU170), and an unclassified *Cryptomonas* (OTU11) (Table S1, 18S OTU table norm.). Comparatively, two unclassified *Eukaryotes* (OTU2 and OTU1), an unclassified *Stramenopiles* (OTU3), an unclassified *Perkinsidae* (OTU13), and an unclassified *Chrysophyceae* (OTU6) had the highest abundance in the > 2.7 μm fraction (Table S1, 18S OTU table norm).

### The Mississippi River Core Microbiome

We defined the core microbiome as those OTUs detectable after normalization in greater than 90% of the sites. The 16S rRNA gene > 2.7 μm and 0.2-2.7 μm core microbiomes consisted of 82 and 98 OTUs, respectively, classified into eight different phyla- Proteobacteria, Actinobacteria, Bacteroidetes, Cyanobacteria, Verrucomicrobia, Chloroflexi, Chlorobi, Gemmatimonadetes- and composed of taxa such as freshwater SAR11 (LD12, Proteobacteria), *Limnohabitans* sp. (Proteobacteria), *Polynucleobacter* sp. (Proteobacteria), acI clade (Actinobacteria), LD28 clade (Proteobacteria), and *Planktothrix* sp. (Cyanobacteria) (Table S1, 16S 0.2-2.7 μm Core Comm. and 16S > 2.7 μm Core Comm.). Core microbome relative abundance in both fractions decreased along the upper river but stabilized in the lower river (Fig. 4A). We confirmed this effect by analyzing the upper and lower core microbiomes separately. Although the total OTU numbers changed (81 and 116 OTUs in the upper MSR and 160 and 144 OTUs in the lower MSR for the > 2.7 μm and 0.2-2.7 μm fractions, respectively), the trends remained the same (Fig. S5).

**Figure 4.**
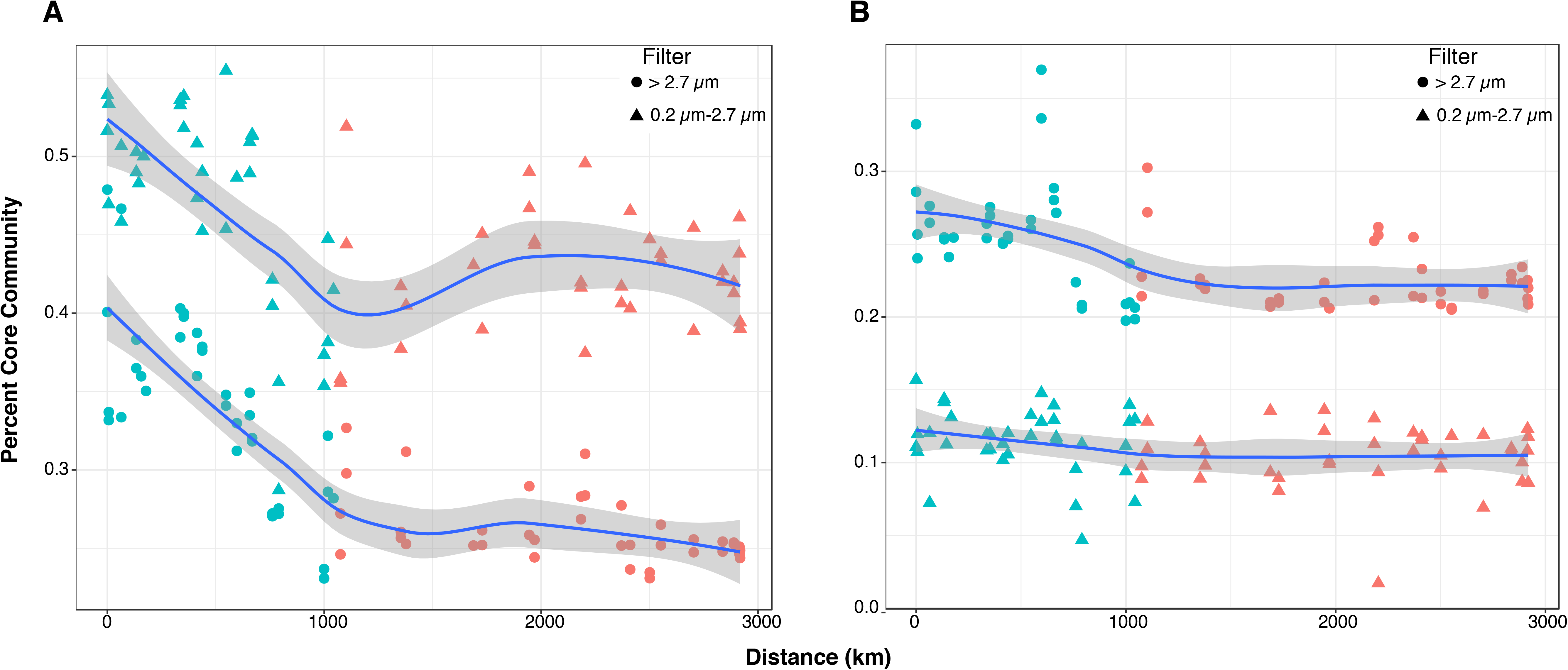
Core microbiome aggregate abundance for the 16S (A) and 18S (B) rRNA gene. In each, triangles and circles points represent 0.2-2.7 μm and > 2.7 μm fractions, respectively. Non-linear regressions with 95% confidence intervals (CI) (gray shading) are provided for reference.

Eighty OTUs comprised the > 2.7 μm 18S rRNA gene core microbiome (Fig. 4B). We classified these as *Alveolata*, *Cryptophceae*, *Nucletmycea*, *Stramenopiles*, or unclassified Eurkaryota (Table S1, 18S > 2.7 μm Core Comm.). Again, consistent with larger organism sizes, and thus fewer OTUs overall, the 0.2-2.7 μm Eukaryotic core microbiome comprised only 21 OTUs (Table S1, 18S 0.2-2.7 μm Core Comm.). These OTUs consisted of *Alveolata*, *Nucletmycea*, *Stramenopiles*, or unclassified Eurkaryota (Table S1, 18S 0.2-2.7 μm Core Comm.). While the 0.2-2.7 μm core microbiome remained relatively stable along the river, the > 2.7 μm core decreased along the upper MSR before stabilizing in the lower river, similarly to that of the prokaryotes (Fig. 4B).

### Network analyses identify taxa associated with and predictive of eutrophication

We applied Weighted Gene Correlation Network Analysis (WGCNA) to identify co-occurring groups of OTUs (submodules) that also had significant associations with the eutrophication nutrients phosphate and nitrate. Of the submodules identified through WGCNA as being most strongly correlated to phosphate and nitrate, we restrict our discussion to those modeled via PLS analysis to predict > 50% of the measured nutrient concentration. Our additional PLS analyses for those submodules predicting < 50% of nitrate and/or phosphate concentrations are included in Figure S6 and Figure S7-S10.

Three submodules in the prokaryotic and eukaryotic fractions were strongly associated with phosphate. The 0.2-2.7 μm prokaryotic submodule most associated with phosphate was composed of 51 OTUs (Table 3), and had moderate correlation between the submodule structure and phosphate (Fig. S7D). Strong correlations between submodule structure and the measured nutrient suggests that individual submodule OTUs that also have strong correlations to the nutrient are the most important organisms associated with that nutrient (Langfelder and Horvath 2008). PLS modeling determined that this prokaryotic submodule predicted 80% of measured phosphate concentrations (Figure S7E, Table 3). Variable importance in the projection (VIP) analysis found OTUs corresponding to an unclassified *Holophagaceae*, an unclassified *Gemmatimonadaceae*, and an unclassified *Burkholderiaceae* were the three most important in the PLS model for phosphate (Fig. 5A, Table S1, 16S 0.2-2.7 μm PO4). OTU322 (Acidobacteria subgroup 6), had moderate correlation to phosphate but high node centrality (n = 51; Fig. 5A, Table S1, 16S 0.2-2.7 μm PO4), corroborating evidence that freshwater sediment Acidobacteria subgroup 6 often occur in co-culture with Alphaproteobacteria and may be metabolically connected (Kielak *et al.* 2016 and refs. within). A submodule from the 0.2-2.7 μm eukaryotic fraction was also highly predictive of phosphate (Table 3, Fig. S8D). We identified the top four VIP taxa as an unclassified *Peronosporomycetes*, an unclassified *Ochrophyta*, an unclassified Eukaryote, and an unclassified *Stramenopiles* (Fig. 6A, Table S1, 18S 0.2-2.7 μm PO4). All four OTUs had Pearson correlations with phosphate greater than 0.60, among which two were negative (Fig. 6A, Table S1, 18S 0.2-2.7 μm PO4). Unfortunately, the most highly interconnected OTU within the submodule remained unclassified even at the Phylum level (Fig. 6A, Table S1, 18S 0.2-2.7 μm PO4). A > 2.7 μm eukaryotic submodule could predict 62% of measured phosphate variance (Table 3). An unclassified Eukaryote and an unclassified *Peronosporomycetes* occupied top two VIP positions (Fig. 6B, Table S1, 16S > 2.7 μm PO4).

**Figure 5.**
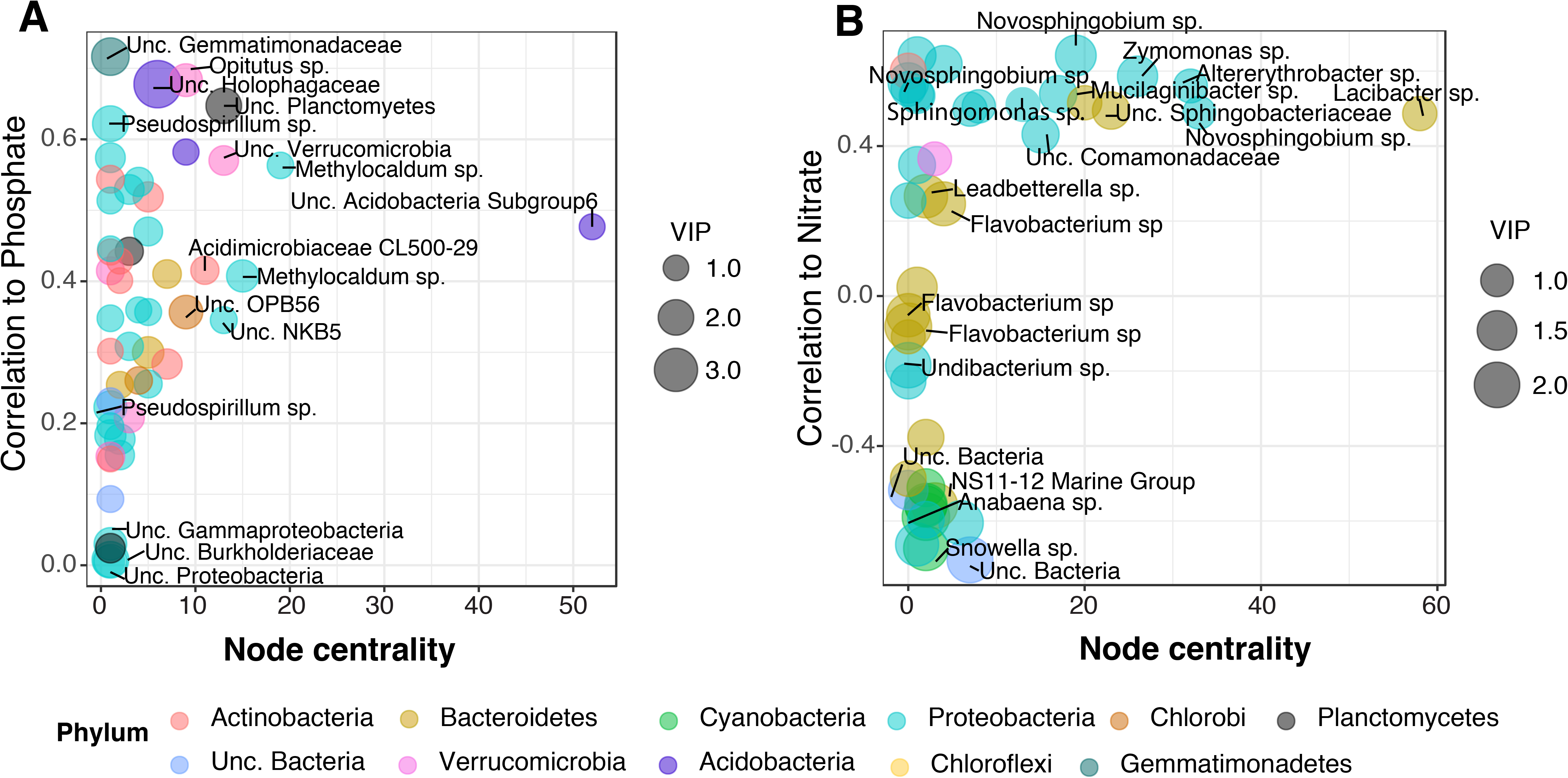
PLS results for the 16S rRNA gene community submodules most associated with phosphate (A) and nitrate (B). OTU correlation with a given nutrient is indicated on the y-axis according to the number of co-correlations (node centrality) on the x-axis. Community fractions: 0.2-2.7 μm (A) and > 2.7 μm (B). Circle size is proportional to VIP scores, with top 10 VIP scoring and top node centrality OTUs labeled with their highest-resolution taxonomic classification and OTU number. Colors represent the taxonomic classification the phylum level. For all OTU taxonomic designations resulting from this analysis, see Table S1 “16S 0.2-2.7μm PO4” and “16S > 2.7μm NO3”.

**Figure 6.**
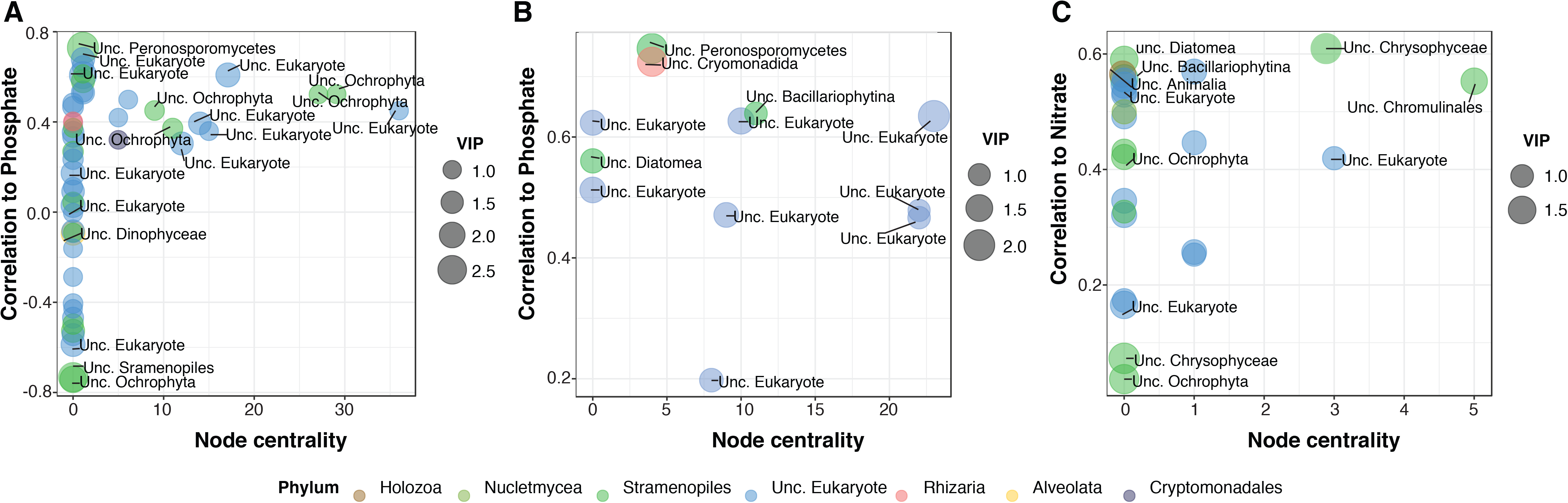
PLS results for the 18S rRNA gene community submodules most associated with phosphate and nitrate. OTU correlation with a given nutrient is indicated on the y-axis according to the number of co-correlations (node centrality) on the x-axis. Community fractions: 0.2-2.7 μm (A) and > 2.7 μm (B, C). Circle size is proportional to VIP scores, with top 10 VIP scoring and top node centrality OTUs labeled with their highest-resolution taxonomic classification and OTU number. Colors represent the taxonomic classification the phylum level. For all OTU taxonomic designations resulting from this analysis, see Table S1 “18S 0.2-2.7μm NO3”, “16S > 2.7μm PO4”, and “16S >2.7μm NO3”

**Table 3.** Correlations between prokaryotic and eukaryotic submodules to nitrate and phosphate, as well as PLS model results

| *Fraction* | *Gene* | *Nutrient* | *Submodule* | *Eigen Correlations* | *Total OTUs* | *PLS model* |
| --- | --- | --- | --- | --- | --- | --- |
| 0.2-2.7 $\mu\text{m}$ | 16S | $\text{NO}_3^{2-}$ | Blue | R = 0.6, P = 7e-08 | 77 | $R^2=0.35$; R = 0.65, P = 1.e-09 |
| > 2.7 $\mu\text{m}$ | 16S | $\text{NO}_3^{2-}$ | Brown | R = 0.56, P = 3e-07 | 133 | $R^2 = 0.69$; R = 0.83, P = < 2.2e-16 |
| 0.2-2.7 $\mu\text{m}$ | 16S | $\text{PO}_4^{3-}$ | Turquoise | R = 0.53, P = 2e-06 | 51 | $R^2=0.80$; R = 0.89, P = < 2.3e-16 |
| > 2.7 $\mu\text{m}$ | 16S | $\text{PO}_4^{3-}$ | Yellow | R = 0.53, P = 1e-06 | 80 | $R^2= 0.48$; R = 0.77, P = 4.88e-15 |
| 0.2-2.7 $\mu\text{m}$ | 18S | $\text{NO}_3^{2-}$ | Green | R = 0.39, P = 6e-04 | 39 | $R^2=0.38$; R = 0.62, P = 2.97e-9 |
| > 2.7 $\mu\text{m}$ | 18S | $\text{NO}_3^{2-}$ | Brown | R = 0.52, P = 2e-06 | 59 | $R^2=0.572$; R = 0.759, P = 6.7e-15 |
| 0.2-2.7 $\mu\text{m}$ | 18S | $\text{PO}_4^{3-}$ | Turquoise | R = 0.56, P = 2e-07 | 56 | $R^2=0.80$; R = 0.89, P = < 2e-16 |
| > 2.7 $\mu\text{m}$ | 18S | $\text{PO}_4^{3-}$ | Red | R = 0.60, P = 2e-08 | 39 | $R^2 = 0.618$; R = 0.799, P = < 2e-16 |

Only two submodules had strong associations with nitrate. The > 2.7 μm prokaryotic submodule most strongly correlated with nitrate contained 133 OTUs (Table 3, Fig. S9A). However, despite a low structural correlation to nitrate, the submodule strongly predicted measured nitrate in the PLS model (Figure S9B, Table 3). The four highest VIP scoring OTUs were an *Anabaena* sp., a *Flavobacterium* sp., an unclassified bacterium, and a member of the *Sphingobacteriales* NS11-12 marine group (Fig. 5B, Table S1, 16S > 2.7 μm NO3). OTUs with the highest node centrality (> 20) belonged to the *Sphingomonadales* (*Alphaproteobacteria*) and *Sphingobacteriales* (Bacteroidetes), and each positively correlated with nitrate (R > 0.48, Fig. 5B, Table S1, 16S > 2.7 μm NO3). The > 2.7 μm eukaryotic submodule most associated with nitrate predicted 57% of observed variation in nitrate concentrations (Table 3). OTUs with the top VIP scores were two unclassified *Chrysophyceae*, an unclassified *Ochrophyta*, and an unclassified Diatom (Fig. 6C, S10; Table S1, 18S > 2.7 μm NO3).

## Discussion

Understanding microbial communities on a whole-river scale is essential for linking the complex relationships between microorganisms, metabolism, and water quality. At 2914 km, this study is the largest transect of the MSR to date, and the first to include both prokaryotic and eukaryotic data by size fractions. This is also the largest river yet surveyed at this level of geographic completion, which may account for some of its distinctive microbiological features. Although the MSR comprised similar aquatic microbial taxa found within other rivers (Zwart *et al.* 2002; Cottrell *et al.* 2005; Ghai *et al.* 2011; Fortunato *et al.* 2013; Jackson *et al.* 2014; Read *et al.* 2015; Savio *et al.* 2015; Meziti *et al.* 2016), it hosted unique relative abundance and core microbiome trends, indicating that rivers have distinguishing microbiome characteristics. Furthermore, we observed two different community regimes separated at the Missouri River confluence and mirrored by the measured physio-chemical properties of the MSR (Fig. 1B). This is an important hydrological feature of the MSR that strongly influences its microbiology, and further distinguishes it from other rivers. Our study also identified taxa associated with, and predictive of, the eutrophication nutrients nitrate and phosphate that provide important targets for future study and may assist in detecting and quantifying imminent changes in river water quality.

MSR microbial communities separated into two distinct fractions of microbial assemblages, 0.2-2.7 μm and > 2.7 μm (ANOSIM, R = 0.65, P = 0.001), throughout the river transect, a common feature of aquatic ecosystems (DeLong *et al.* 1993; Crump *et al.* 1999; Allgaier and Grossart 2006; D’Ambrosio *et al.* 2014; Jackson *et al.* 2014). However, despite sharing similar taxa with other rivers (Ghai *et al.* 2011; Fortunato *et al.* 2012; Crump *et al.* 2012; Read *et al.* 2015; Savio *et al.* 2015; Meziti *et al.* 2016; Niño-García *et al.* 2016), MSR microbial community abundances were distinct at a phylum level from previous work on Thames River (Read *et al.* 2015), Danube River (Savio *et al.* 2015), Canadian Boral Rivers (Niño-García *et al.* 2016), Toolik catchment (Crump *et al.* 2012), and Yenisei River (Kolmakova *et al.* 2014). Specifically, within the 0.2-2.7μm MSR prokaryotic fraction, Proteobacteria remained the most abundant phylum throughout the river, while Bacteroidetes and Actinobacteria decreased downriver (Fig. 3). In contrast, other river studies found headwaters dominated by Bacteroidetes that gave way to more abundant Actinobacteria and Proteobacteria further downriver (Read *et al.* 2015; Savio *et al.* 2015). Similarly, the trends observed in the upper portion of the MSR (Staley *et al.* 2013), where Bacteroidetes and Actinobacteria decreased while Proteobacteria increased, diverged from the trends found in this study (Fig. 3). Further, while no clear increasing or decreasing trends were observed in Payne *et al.* (2017), Actinobacteria occurred with the highest relative abundances in the planktonic fraction, whereas in this study, Proteobacteria were the most abundant phylum. This may be attributable to the difference in sampling year or possibly methodology, such as an amplification step that was not undertaken in this work (Payne *et al.* 2017). Furthermore, although many of the dominant taxa found in other rivers also occupied the MSR (LD12 (freshwater SAR11, *Alphaproteobacteria*), *Limnohabitans* sp. (*Betaproteobacteria*), LD28 (*Betaproteobacteria*), acI clade (Actinobacteria), and *Algoriphagus* sp. (Bacteriodetes) (Ghai *et al.* 2011; Fortunato *et al.* 2013; Kolmakova *et al.* 2014; Read *et al.* 2015; Savio *et al.* 2015; Meziti *et al.* 2016), we observed OTU-level abundance differences at a whole-river scale. For example, OTUs belonging to the acI and LD12 freshwater clades continually increased in abundance towards the river mouth in the Thames and Danube Rivers (Read *et al.* 2015; Savio *et al.* 2015), whereas MSR OTUs classified as acI (OTUs 3, 4 and 7) and LD12 (OTU 11) did not show any distinct change in abundance throughout the transect. Specific to studies of portions of the MSR.

The generally greater richness (Table S1, Richness comparisons) and decreased core microbial community abundances (Fig. 4) in the lower vs. upper MSR contrasted predictions by the RCC (Vannote *et al*., 1980) and observations in previous studies from both large and small rivers (Read *et al.* 2015; Savio *et al.* 2015, Niño-García *et al.* 2016) where these trends were reversed. Importantly, we did not sample the true headwaters of the MSR (Lake Itasca to above St. Cloud), and at the point of first sampling, the MSR already constituted an eighth order river. Therefore, some of the trends predicted by the RCC may have been missed, although increasing richness was also observed in the lower MSR during a different year (Payne *et al.* 2017).

Ultimately, the variant observations between this MSR survey and other river studies may result from different sampling methodologies or the particular timing of sampling. Previous studies conducted on six artic rivers (Crump *et al.* 2009), two smaller temperate rivers (Parker and Ipswich Rivers) (Crump and Hobbie 2005), and the upper portion of the MSR (Staley *et al.* 2015) showed that seasonality was important in structuring bacterial communities over three year and two year sampling periods.

However, our observed differences may also stem from biological signal related to unique environmental conditions, human impacts (Meziti *et al.* 2016; Payne *et al.* 2017), and changes in hydrology and the level of river engineering (Ruiz-González *et al.* 2013, 2015; Freimann *et al.* 2015; Niño-García *et al.* 2016) with distance. For instance, the most striking difference between the MSR and other rivers was the distinct separation of prokaryotic and eukaryotic community regimes at the Missouri River confluence. This separation matched physio-chemical changes observed for the MSR such as increased Strahler river order (8 to 10) and more consistent nutrient concentrations in the lower river compared to the upper (Fig. 1B) (Pierson *et al.* 2008). These results are bolstered by another observation depicting a major separation of communities above and below the Missouri confluence, observed during a different year (2012) and roughly three months prior (late summer) to the current study (Payne *et al.* 2017). This similarity to our results suggests the major influence of the Missouri River confluence is not seasonally dependent.

Although the overall microbial community patterns in the MSR differed from other rivers, we hypothesize that similar ecological processes drove the separation between the upper and lower MSR communities, namely changes in the importance of immigration, emigration, and resource gradient dynamics (Crump *et al.* 2012; Savio *et al.* 2015; Niño-García *et al.* 2016).

Specifically, our data suggest that mass effects play a role in structuring microbial communities in the upper MSR, although instead of only in the headwaters, this process continues for almost a third of the length of the river, possibly due to contributions from large tributaries and abundant dams. This contrasts with the conclusion made by Staley *et al.* (2015) for a smaller transect in the upper MSR, where they suggested species sorting was the primary influence structuring upper MSR microbial communities. Increased turbidity correlated with decreased core microbiome relative abundance (Spearman rank correlation, > 2.7 μm R = 0.53; 0.2-2.7 μm R = 0.63) in the upper MSR, and nutrients like phosphate and nitrate continually increased along the upper river in the current study (Fig. 1B). Similarly, richness also increased along the upper river (except in the 0.2-2.7 μm 18S communities) (Fig. S3). These patterns are consistent with communities under the influence of strong mass effects in the upper MSR, whereby inputs from allochthonous sources continually contributed nutrients, particulate matter, and additional microbial taxa.

Conversely, nitrate, phosphate, turbidity, and core microbiome relative abundance all stabilized in the lower MSR (Figs. 1B, 4), suggesting a change in the relative influence of external sources for these variables. We speculate that once the MSR grew to a tenth order river, its large volume and size buffered it from allochthonous inputs. Nevertheless, overall richness was greater in the lower river compared to the upper (except in the 0.2-2.7 μm 18S communities) (Table S1, Richness comparisons), and, like the upper river, richness generally increased in the lower river (Fig. S3), matching a previous observation (Payne *et al.* 2017). However, since the relative abundance of core microbiome taxa remained comparatively stable in the lower river (Fig. 4), increased richness likely occurred primarily from the emergence of low abundance OTUs. The extent to which a local or regional event impacts downriver populations is dependent on the success of allochthonous taxa associated with such an event to become established within the autochthonous population (Crump *et al.* 2012; Niño-García *et al.* 2016). The larger the river, the less likely any newly immigrated taxa will establish dominance. Thus, a plausible scenario for the regime change observed at the Missouri River confluence is that the influence of mass effects was drastically reduced by the increased size of the lower river, and that allochtonous taxa immigrating there were relegated to lower abundances.

Alternatively, some of the increase in richness may have occurred through native community differentiation within the lower river. An overall community shift from a mixture of allochthonous members to a “native” population requires growth rates greater than residence time over a given distance (Crump *et al.* 2004). Though river speed increased along the MSR, the effective residence time also increased since taxa no longer experienced rapidly changing environmental variables (Fig. 1B). As a result, the lower river may have provided opportunities for microbial community differentiation based on microniches within the river, which is possible considering average prokaryotic growth rates (Savio *et al.* 2015), especially among particle-associated (> 2.7 μm) taxa (Crump *et al.* 1998). Thus, although the aggregate patterns in particular phyla and the overall taxonomic richness within the MSR differ from other systems, similar ecological processes may still drive these patterns, but the relative proportion of the river whereby mass effects vs. species sorting dominates fosters unique community assemblages (Niño-García *et al.* 2016). To resolve the relative importance of mass effects vs. species sorting in the upper and lower rivers, future work will need to incorporate explicit measurements of immigrating and emigrating taxa, as well as growth rates for different organisms at different geographic positions.

Within this dynamic river system, we also sought to define the microorganisms most associated with eutrophication nutrients to provide specific taxa for future study of nitrate and phosphate uptake and/or metabolism, and also as plausible biological indicators of river trophic state. By focusing on OTUs with the highest VIP scores in the PLS models, we identified several bacterial and eukaryotic taxa that fit these criteria (Figs. 5, 6). Organisms in the *Sphingomonadaceae* (e.g., *Novosphingobium* spp.) contributed strongly to the PLS models predicting nitrate with both > 2.7 μm (Fig. 5B) and 0.2-2.7 μm (Fig. S6A) size fractions (Table S1, 16S 0.2-2.7 μm VIP NO3 and 16S > 2.7 μm VIP NO3). *Novosphingobium* spp. isolates have previously been associated with eutrophic environments (Trusova and Gladyshev 2002; Zwart *et al.* 2002; Addison *et al.* 2007; Li *et al.* 2012) and some can reduce nitrate (Addison *et al.* 2007; Li *et al.* 2012), making these specific OTUs candidates for nitrate metabolism in the river water column. An *Anabaena* sp. (OTU40) from the core microbiome (Table S1, 16S > 2.7 μm Core Comm.) had the top VIP score within the > 2.7 μm submodule that could predict 69% of nitrate concentrations, and correlated negatively with nitrate (Table 3, Fig. 5B). The nitrogen-fixing *Anabaena* spp. (Allen and Arnon 1955) typically bloom in low dissolved inorganic nitrogen (DIN) conditions (Wood *et al.* 2010), making the absence of these consistent with high DIN. An unclassified *Holophagaceae* OTU (OTU33) had the highest VIP score within the prokaryotic 0.2-2.7 μm submodule that predicted 80% of the variance in phosphate concentrations (Table 3, Fig. 5A). This same OTU was also a key driver of the beta diversity separation between upper and lower river communities (Table 2), and a member of the core microbiome (Table S1, 16S 0.2-2.7 μm Core Comm.). The *Holophagaceae* belong to the Acidobacteria phylum, and the Ohio River contained a much higher abundance of Acidobacteria relative to other tributaries in a previous study (Jackson *et al.* 2014). Notably, Acidobacteria increased with river distance in our study as well (Fig. S4), with a peak in the 0.2-2.7 μm fraction near the Arkansas River confluence. This increase in “free-living” Acidobacteria downriver is distinct from other whole river studies, making these organisms, and the *Holophagaceae* OTUs in particular, potentially important organisms for the MSR river basin specifically.

Within Eukaryotes, multiple different algae, diatom, and Oomycetes OTUs occupied submodules highly predictive of nitrate and phosphate (Fig. 6A-C), and specifically *Chrysophyceae* taxa from both size fractions correlated strongly with nitrate (Fig. 6C, Fig. S6E). *Chrysophyceae* (golden algae) commonly occupy river systems (Necchi Jr 2016) including the MSR (Korajkic *et al.* 2015), can be autotrophic and mixotrophic (Jansson *et al.* 1996), and may serve as predators of prokaryotes (Caron *et al.* 1990). Further, multiple OTUs classified as *Peronosporomycetes* were important in predicting phosphate in both size fractions (Fig 6A, B, Table S1, 18S 0.2-2.7 μm VIP PO4 and 18S > 2.7 μm VIP PO4). *Peronosporomycetes* are fungus-like eukaryotic organisms known to be pathogenic in fish, plants, and mammals (Dick 2003; Islam and von Tiedemann 2011). While we also identified many other eukaryotic OTUs as important predictors of nutrients, poor taxonomic resolution hindered our ability to discuss them further. Improved cultivation and systematics of key microbial eukaryotes will be vital to understanding river nutrient dynamics.

While the most geographically comprehensive analysis to date for the MSR, this study only encompasses a snapshot in time. As noted previously, seasonal changes that have been observed in other rivers (Crump and Hobbie 2005; Crump *et al.* 2009; Smith *et al.* 2010; Fortunato *et al.* 2013; Staley *et al.* 2015) undoubtedly influence this dynamic system, though the permanence of the regime change at the Missouri River confluence remains in question. Future studies should incorporate microbial responses, at a whole-river scale, to seasonal pulse events (e.g. rain, snow melt), and how river size and volume may buffer local microbial communities from allochthonous inputs (Zeglin 2015). Our current research highlights the distinctiveness of MSR microbial communities and the complexities influencing their structure within the MSR ecosystem. The observed association between changes in Strahler’s river order, nutrient dynamics, and community composition indicates the importance of hydrology on the spatial dynamics structuring microbial communities (Freimann *et al.* 2015; Zeglin 2015; Niño-García *et al.* 2016) and provides baseline information for future MSR studies that incorporate greater temporal and spatial resolution. With river water quality of growing local and global importance (Vorosmarty *et al.* 2010; Russell and Weller 2013), the candidate taxa predictive of eutrophic nutrients determined herein also provide important targets for further research into their roles for indicating, and potentially improving, river health.

## Acknowledgements

This work was supported by the Louisiana State University Department of Biological Sciences, College of Science, and Office of Research and Economic Development; and the College of the Environment at the University of Washington. Portions of this research were conducted with high performance computing resources provided by Louisiana State University (http://www.hpc.lsu.edu). The authors thank Dr. Caroline Fortunato and Dr. Gary King for thoughtful comments on the manuscript. The authors also thank the countless volunteers, schools, and organizations that facilitated the research. We specifically thank Pete Weess, Jessica Zimmerman, Katy Welch, Brian Moffitt, and David Cheney for helping organize the shipment of coolers between sites, and the OAR Northwest sponsors- Seattle Yacht Club Foundation, Yeti Coolers, and the National Mississippi River Museum and Aquarium. A full list of sponsors and volunteers can be found on the OAR Northwest website (oarnorthwest.org). We would also like to thank Dr. Matthew Sullivan and Dr. Simon Roux for their help with coding the WGCNA and sPLS analyses. Lastly, we would like to thank Mrs. Ginger Thrash, who connected the Thrash lab to OAR Northwest.

## Supplemental Figure Legends

**Figure S1.** NMDS results for the 16S (A) and 18S (B) rRNA gene communities. In each, circles and triangles represent the > 2.7 μm and 0.2-2.7 μm fractions, respectively.

**Figure S2.** Hclust results for the 16S (A) and 18S (B) rRNA gene communities.

**Figure S3.** Separated upper and lower MSR Richness and Evenness indexes for the 16S > 2.7 μm (A) and 0.2-2.7 μm (B) and 18S > 2.7 (C) and 0.2-2.7 μm (D) communities. Linear regressions with 95% CI (gray shading) are provided for reference.

**Figure S4.** Relative abundance, by phylum, according to transect distance, for phyla accounting for > 0.1% of the total reads for the 16S rRNA (A, B) and 18S rRNA genes (C, D) > 2.7 μm (A, C) and 0.2-2.7 μm (B, D) communities. Non-linear regressions with 95% CI (gray shading) are provided for reference.

**Figure S5.** Separated upper and lower MSR core microbiome aggregate abundance for the 16S rRNA gene communities. For the upper and lower river, the core microbiome was defined separately requiring OTUs to have greater than one read in 90% of the samples. > 2.7 μm (A) and 0.2-2.7 μm (B) 16S rRNA gene communities in the upper and lower MSR.

**Figure S6.** PLS results for the 0.2-2.7 μm and > 2.7 μm 16S (A-D) and 18S (E-H) rRNA gene community for selected submodules with nitrate and phosphate and a VIP score > 1. Correlation of submodule OTUs to nitrate and phosphate according to the number of co-correlations (node centrality) for 0.2-2.7 μm (A, B, E, F) and > 2.7 μm (C, D, G, H) 16S and 18S rRNA gene communities. Circle size is proportional to VIP scores, with top 10 VIP scoring and top node centrality OTUs labeled with their highest-resolution taxonomic classification and OTU number. Colors represent the taxonomic classification the phylum level.

**Figure S7.** WCGNA results for 0.2-2.7 μm 16S rRNA gene community submodules of interest based on Pearson correlations to environmental measurements (A). Color shading depicts the strength of the Pearson correlation with individual submodule’s eigenvalue. Boxes indicate the selected submodule with its Pearson correlation and P-value. Graphs of the relationship between the selected submodule OTUs and the strength of the individual OTUs to nitrate (B) and phosphate (D). PLS regression of the predicted nutrient concentrations versus measured nutrient concentrations (C, E) using the corresponding selected submodules. Linear regressions with 95% CI (gray shading) are provided for reference.

**Figure S8.** WCGNA results for 0.2-2.7 μm 18S rRNA gene community submodules of interest based on Pearson correlations to environmental measurements (A). Color shading depicts the strength of the Pearson correlation with individual submodule’s eigenvalue. Boxes indicate the selected submodule with its Pearson correlation and P-value. Graphs of the relationship between the selected submodule OTUs and the strength of the individual OTUs to nitrate (B) and phosphate (D). PLS regression of the predicted nutrient concentrations versus measured nutrient concentrations (C, E) using the corresponding selected submodules. Linear regressions with 95% CI (gray shading) are provided for reference.

**Figure S9.** WCGNA results for > 2.7 μm 16S rRNA gene community submodules of interest based on Pearson correlations to environmental measurements (A). Color shading depicts the strength of the Pearson correlation with individual submodule’s eigenvalue. Boxes indicate the selected submodule with its Pearson correlation and P-value. Graphs of the relationship between the selected submodule OTUs and the strength of the individual OTUs to nitrate (B) and phosphate (D). PLS regression of the predicted nutrient concentrations versus measured nutrient concentrations (C, E) using the corresponding selected submodules. Linear regressions with 95% CI (gray shading) are provided for reference.

**Figure S10.** WCGNA results for > 2.7 μm 18S rRNA gene community submodules of interest based on Pearson correlations to environmental measurements (A). Color shading depicts the strength of the Pearson correlation with individual submodule’s eigenvalue. Boxes indicate the selected submodule with its Pearson correlation and P-value. Graphs of the relationship between the selected submodule OTUs and the strength of the individual OTUs to nitrate (B) and phosphate (D). PLS regression of the predicted nutrient concentrations versus measured nutrient concentrations (C, E) using the corresponding selected submodules. Linear regressions with 95% CI (gray shading) are provided for reference.

## Supporting Tables

Supplemental Table S1 is a spreadsheet, TableS1.xlsx. Tabs include General Information, Spearman Rank, 0.2-2.7 μm and > 2.7 μm 16S and 18S rRNA gene Core communities, 0.2-2.7 μm and > 2.7 μm 16S and 18S rRNA WGCNA, and 16S and 18S rRNA gene OTU tables (trimmed and normalized).

Additional Supplemental Information, including R scripts, Table S1, and our Mothur workflow are hosted on the Thrash Lab website at: thethrashlab.com/publications.

